# The key miRNAs for metastasis and survival of colon adenocarcinoma

**DOI:** 10.1101/757617

**Authors:** Mengxi Yu, Wei Li, Chuang Zhang, Feng Xie, Chengyan He

## Abstract

WGCNA (weighted gene co-expression network analysis) can provide a system-level insight into molecular interaction. To understand the co-expression pattern of miRNA in the progress of colon cancer development, this study constructed the WGCNA co-expression network to find the key microRNA of clinical significance and therefor provide possible personalized treatment targets. The miRNA expression data and clinical follow-up information of 442 valid colon cancer samples were obtained from TCGA. After the co-expression network was constructed, we found the turquoise module has the strongest correlation with stage IV. The miRNAs of the key module are enriched by GO and analyzed by KEGG, and the HUB-miRNAs are screened out through the network node degree. For clinical significance validation, data of 65 cases of colon cancer in GES29623 were analyzed by Cox regression and Kaplan-Meier, in 9 HUB-miRNAs, 2 miRNAs associated with overall survival rate were screened out. They can divide patients into high-risk group and low-risk group. Conclusion: has-miR-126-3p and has-miR-137 are the key miRNAs of COAD metastasis, they are of prognostic value in COAD.

## 1. Introduction

Colorectal cancer (CRC) is one of the most common cancers in the world and the fourth most common cause of cancer-related death(1, 2). The molecular mechanism of colon cancer has yet to be fully discussed. The survival rate of colon cancer patients varies significantly depending on the stage at the time of discovery. Early diagnosis is a key factor for patients with colorectal cancer to achieve a better prognosis(3). The survival rate of early colon cancer (I) was about 90%, while the survival rate of advanced colon cancer (Ⅲ) decreased to 65%, due to the lack of sensitive and effective biomarkers, the 5-year survival rate of patients with distant metastasis decreased to 12.5%(4). Early screening has been proven to have a great impact on increasing survival rate(5). To understand the mechanism of the occurrence development invasion and metastasis of colon cancer is essential to develop targeted strategies.

MiRNA is a small non-coding RNA, that regulates gene expression and participates in almost all biological processes of the human body(6). They can affect the occurrence and prognosis of diseases by regulating mRNA in target cells, which is the regulator of a variety of human diseases(7). Functional studies have shown that all known tumor processes, including apoptosis, proliferation, survival and metastasis, are regulated by miRNA(8). In recent years, the key role of miRNA in the tumor has attracted wide attention(9). As a biomarker of many kinds of cancer, miRNA has a broad application prospect. The unique ability of miRNA to affect multiple downstream pathways represents a new approach to cancer treatment(10). Several kinds of miRNA as biomimetic tumor inhibitors have begun the first phase of clinical trials(11). At present, there are still few studies about the role of miRNA in COAD (colon adenocarcinoma), and the current studies are mostly based on the comparison of differences between groups, limited by the number of samples these studies can only be analyzed within a very limited statistical framework.

WGCNA (weighted gene co-expression network analysis) is a systematic biological analysis strategy that can be used to analyze gene association patterns between a large number of samples and identify highly collaborative gene sets without information loss. The key biomarkers or therapeutic targets can be found according to the inherent association between gene sets and phenotypic correlation(12). Compared with other analysis tools, WGCNA can look for gene sets in massive gene information, it makes full use of the information of each sample, and eliminates the problem of multiple hypothesis test correction. It has played an important role in the field of biomarker discovery in recent years. It has been reported that WGCNA as an analyze tool has been applied to a variety of cancers, such as oral cancer pancreatic ductal adenocarcinoma and gastric cancer, to study the relationship between gene expression matrix and clinical characteristics, in order to determine the rules for predicting the survival outcome of patients(13-15).

## 2. Materials and methods

### 2.1 Samples and preprocessing

The miRNA sequencing and clinical data of 455 COAD samples were obtained from TCGA database (https://cancergenome.nih.gov/). Among the related clinical follow-up data, the total survival time (days), staging, lymphatic metastasis, body weight and condition were selected for follow-up analysis. The gene expression profile of GSE S29623 for validation was downloaded from Gene Expression Omnibus (GEO) database (http://www.ncbi.nlm.nih.gov/geo/). The further data processing was performed on the RStudio (v1.1.462) and complied with the human subject protection and data access policies of TCGA and GEO.

### 2.2 Weighted gene co-expression analysis

A goodSamplesGenes function is used to test whether the miRNA data matrix is qualified. After data cleaning a co-expression network is constructed according to the protocol of WGCNA in R environment. According to the principle of WGCNA analysis, the similarity of different gene expression profiles is understood based on the Pearson correlation coefficient, which is an index to measure the consistency of gene expression profiles among samples(16). Then, the similarity matrix is transformed into the adjacent matrix by using the power adjacency function, and the connection strength between the pairs of nodes is encoded. The selected soft threshold function is used to evaluate whether the topology of the network is scale-free or not. β is a soft threshold-power parameter. We went through the parameters and determined that the power with β = 2 (scale-free R2 ≥ 0.87) can ensure the scale-free network (Fig 1). Then, hierarchical clustering trees are constructed from different branches of trees that represent different gene modules. According to different needs, there are three different ways to build the network and identify the module. In this study, we use the one-step method function for network construction and consistency module detection.

**Figure 1.**
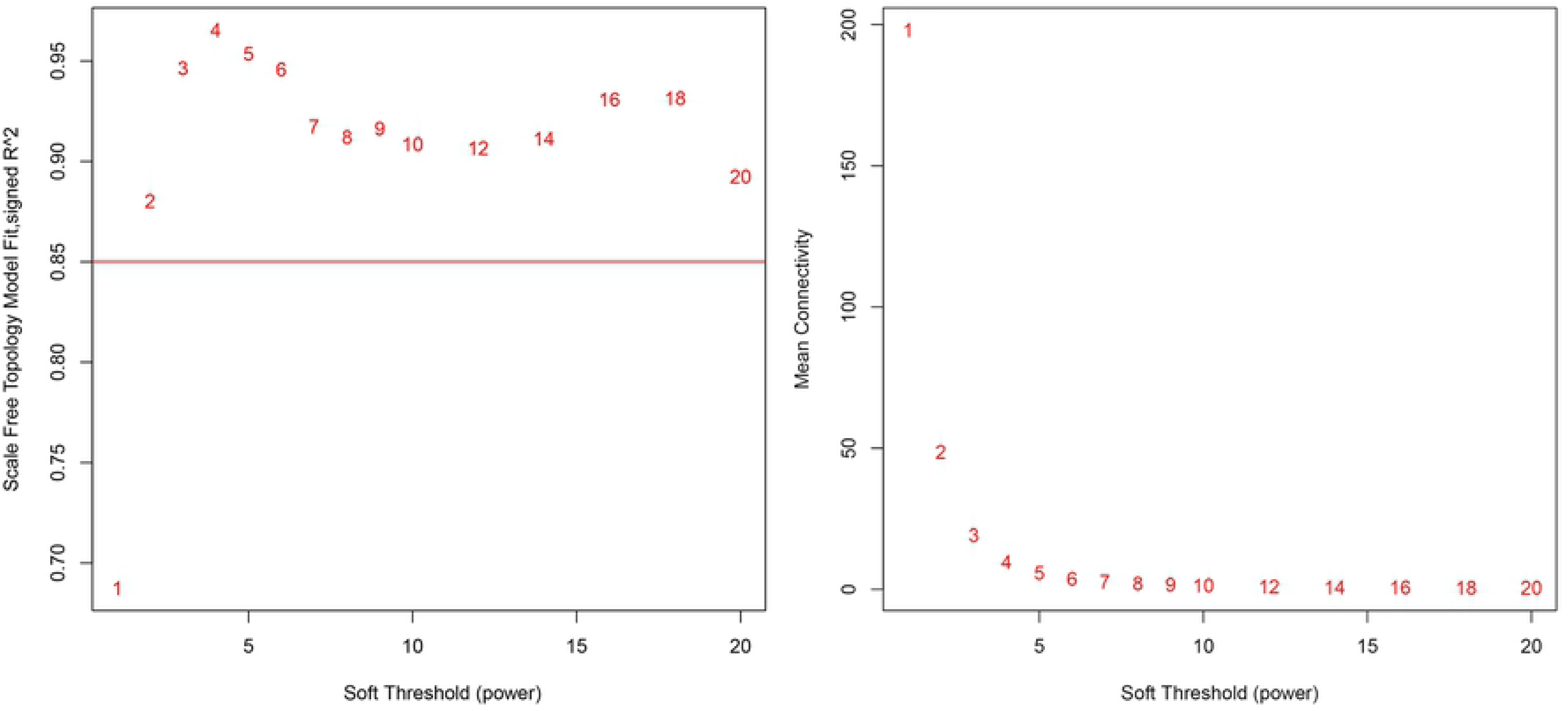
Power. The power with β = 2 (scale-free R2 ≥ 0.87) can ensure the scale-free network.

### 2.3 Identifying important modules and key miRNAs related to target clinical features

We represent each module by calculating the ME (module eigengene), which can be used to capture the largest variation in the module to identify meaningful modules. We associate ME with clinical features to quantify module-feature correlation and find the most important association. Through the signed KME and module Eigengenes function, and using miRNet to build a network, through node degree, we screened out the key miRNAs in the relevant modules (17).

### 2.4 Functional analysis

A large number of target genes were introduced through mirPathV3 database, and a variety of gene analysis tools were used to analyze the modules of interest, miRNA and target genes, and the information of cell composition, molecular function and KEGG pathway were obtained(18).

### 2.5 Hub-miRNAs Verification

In order to ensure the reliability of the results, it is necessary to ensure that the data used for verification come from different samples, so we use databases from different sources to carry out comprehensive survival analysis and evaluation of the key miRNA. We used GSE29623 (N = 65), a colon cancer-related miRNA chip from the GEO database, to analyze the survival of the screened miRNA by cox regression and the KM method. Finally, the HUB-miRNA is verified again by Oncomir (http://www.oncomir.org/).

## 3 Results

3.1 The clinical information and miRNA sequencing of 442 samples were screened out by goodSamplesGenes function, each sample contains 2149 MIRNA expression information, and finally, a matrix containing 442 × 2149, a total of 949858 miRNA is obtained. WGCNA analysis showed that there were 9 functional modules (black, green, magenta, red, yellow, pink, turquoise, blue, brown). The grey model represents genes that cannot be classified and have nothing to do with the disease being analyzed (Fig 2). The miRNAs selected in each category draws a network heat map in its corresponding module (Fig 3) The hierarchical clustering tree was obtained (Fig 4). Each module has a high degree of independence. Each module was combined with their corresponding clinical traits. According to the combination, we found the module with the strongest correlation with stage IV, the phase with distant metastasis, is the turquoise module, P < 0.003 (Fig 5).

**Figure 2.**
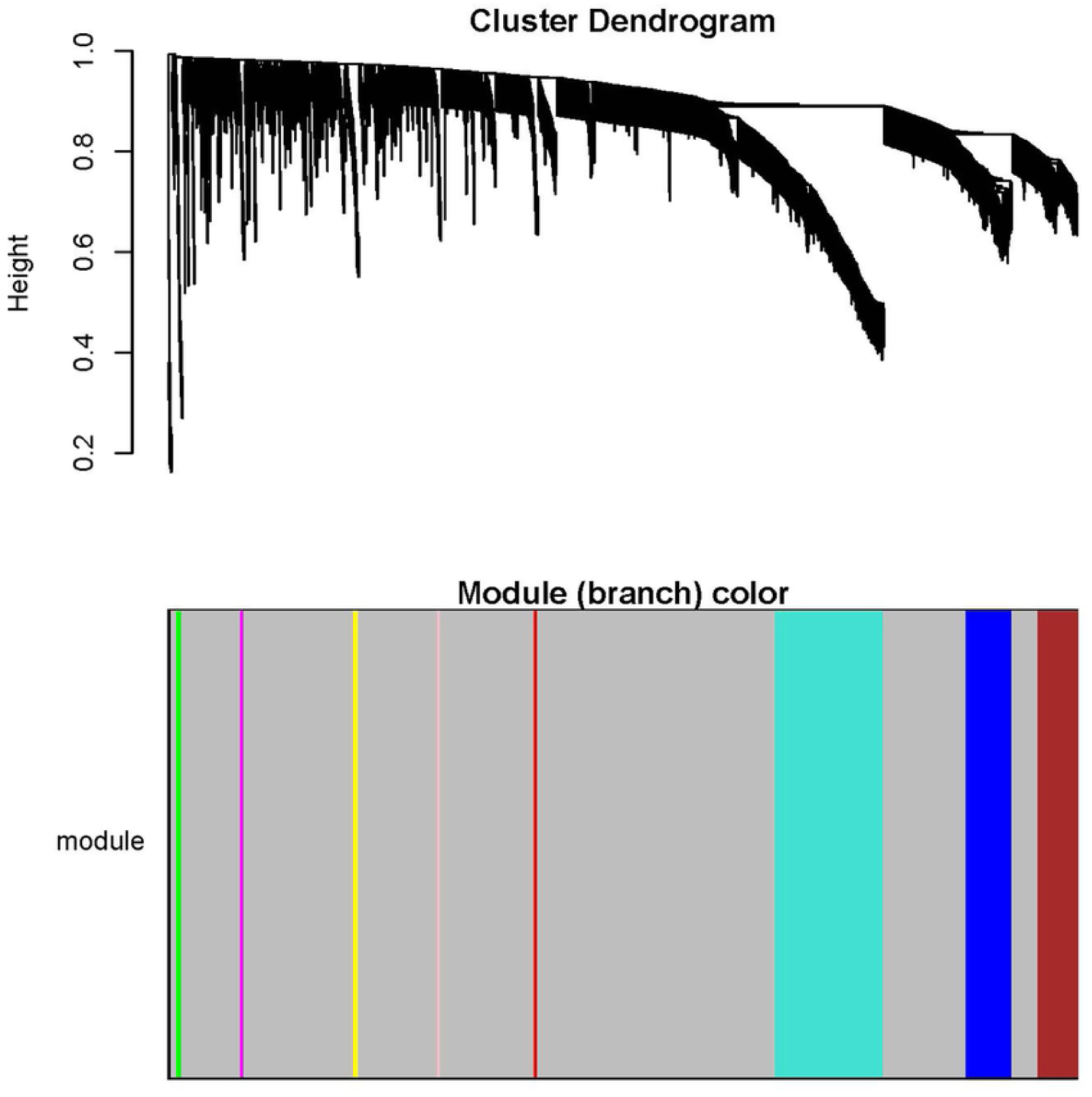
Modules. Nine functional modules (black, green, magenta, red, yellow, pink, turquoise, blue, brown) were obtained after scale free network construction.

**Figure 3.**
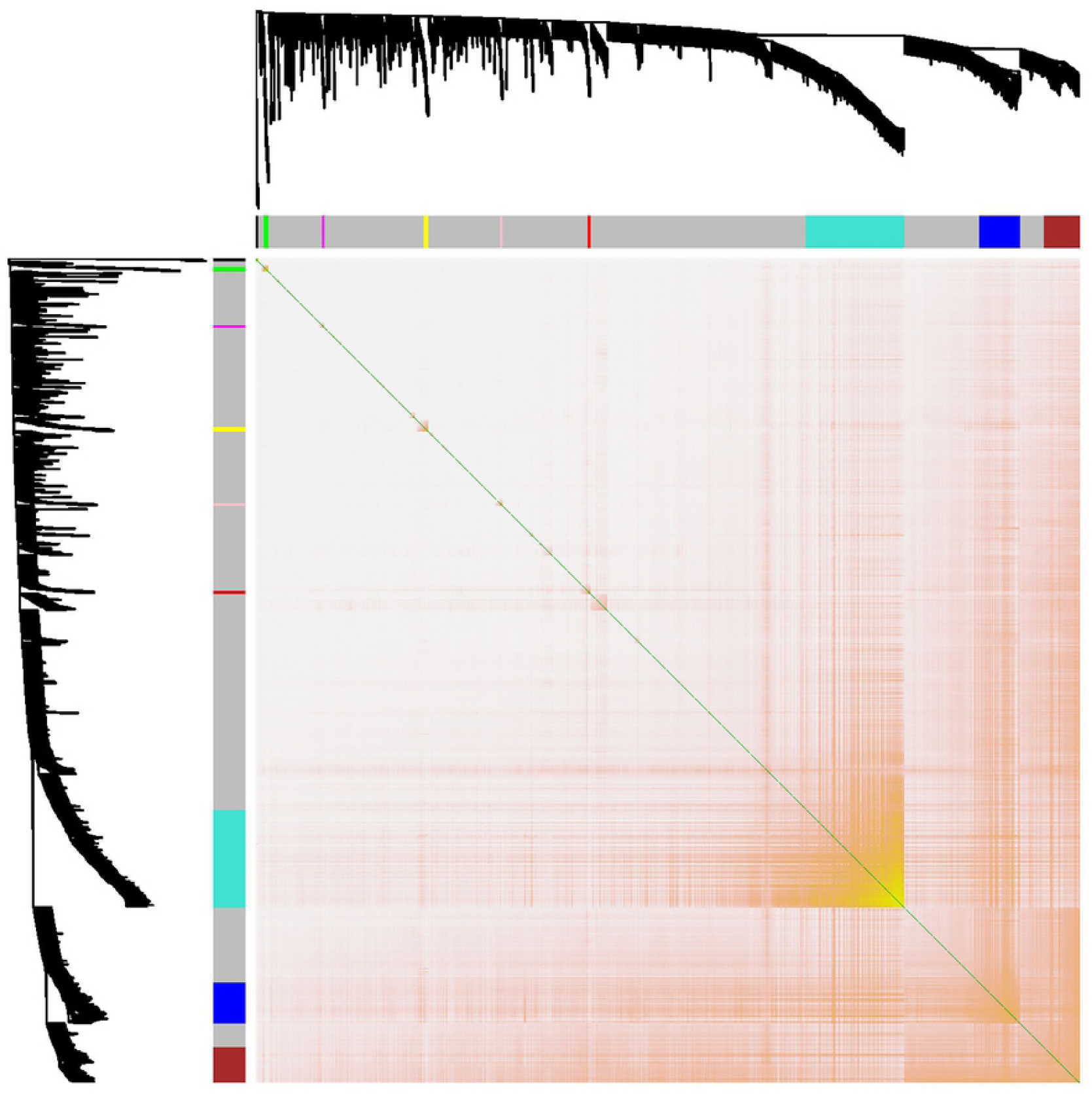
The network heat map. The network heat map is plotted.

**Figure 4.**
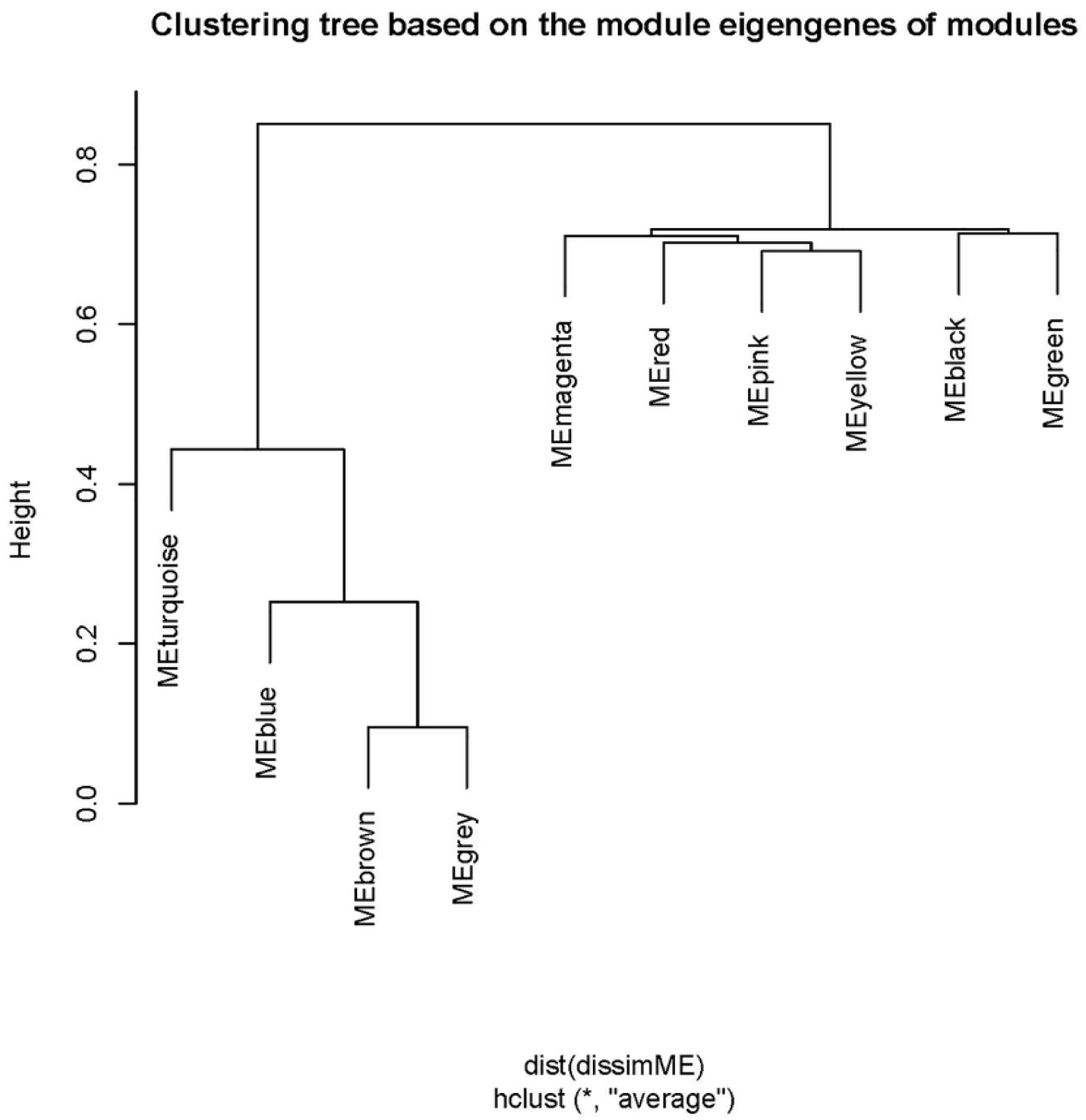
The hierarchical clustering tree. The hierarchical cluster analysis of 10 modules.

**Figure 5.**
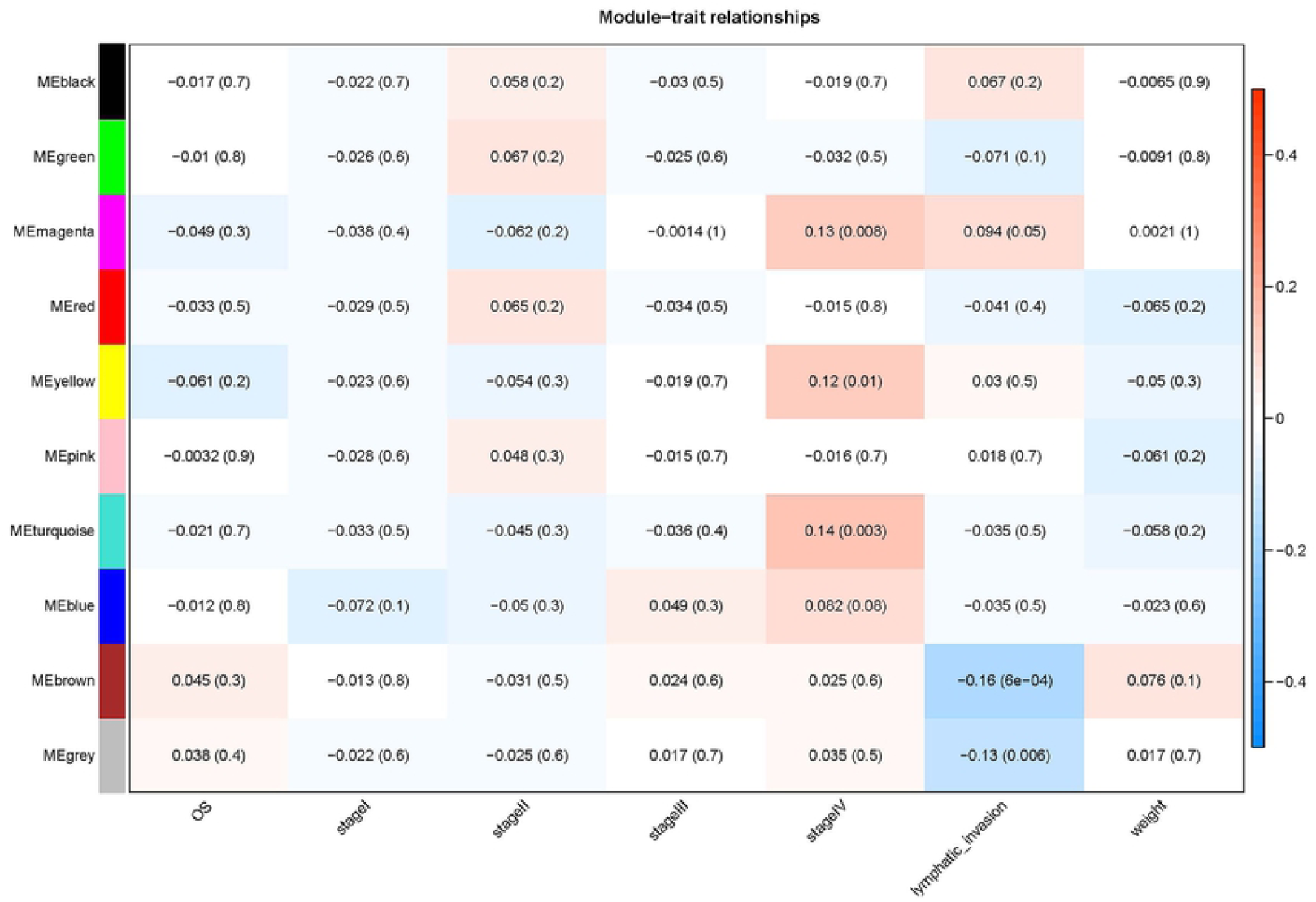
Module-clinic. The turquoise module had the strongest correlation with distant metastasis.

### 3.2 Extraction of key miRNAs for functional annotation KEGG aathway enrichment analysis

In order to functionally annotate the center miRNA, we analyzed the miRNAs using miRNet, to determine the KEGG and GO terms. According to the functional analysis of KEGG and GO, the turquoise module is mainly concentrated in cancer-related words such as proteoglycans in cancer and TGF-beta signals. There are 85 miRNAs in the turquoise module that meet the condition of | kME | > = threshold (0.8), to narrow the range, we set up the interaction network of miRNA-GENE by using all 255 miRNA in the turquoise module and their target genes verified by experiments in intestines (19). 9 miRNAs in the center of the network screened out by node degree as the HUB-miRNAs, hsa-mir-186-5p, hsa-mir-27b-3p, hsa-mir-130a-3p, hsa-mir-137, hsa-mir-494-3p, hsa-mir-337-3p, hsa-mir-31-3p, Hsa-mir-126-3p, hsa-mir-29c-5p (Fig 6). Through KEGG pathway REACTOME and GO analysis of key miRNA, it can be found that these 9 candidate miRNAs are associated with a variety of cancers such as colon cancer, prostate cancer, renal cell carcinoma, etc(Fig 7-10).

**Figure 6.**
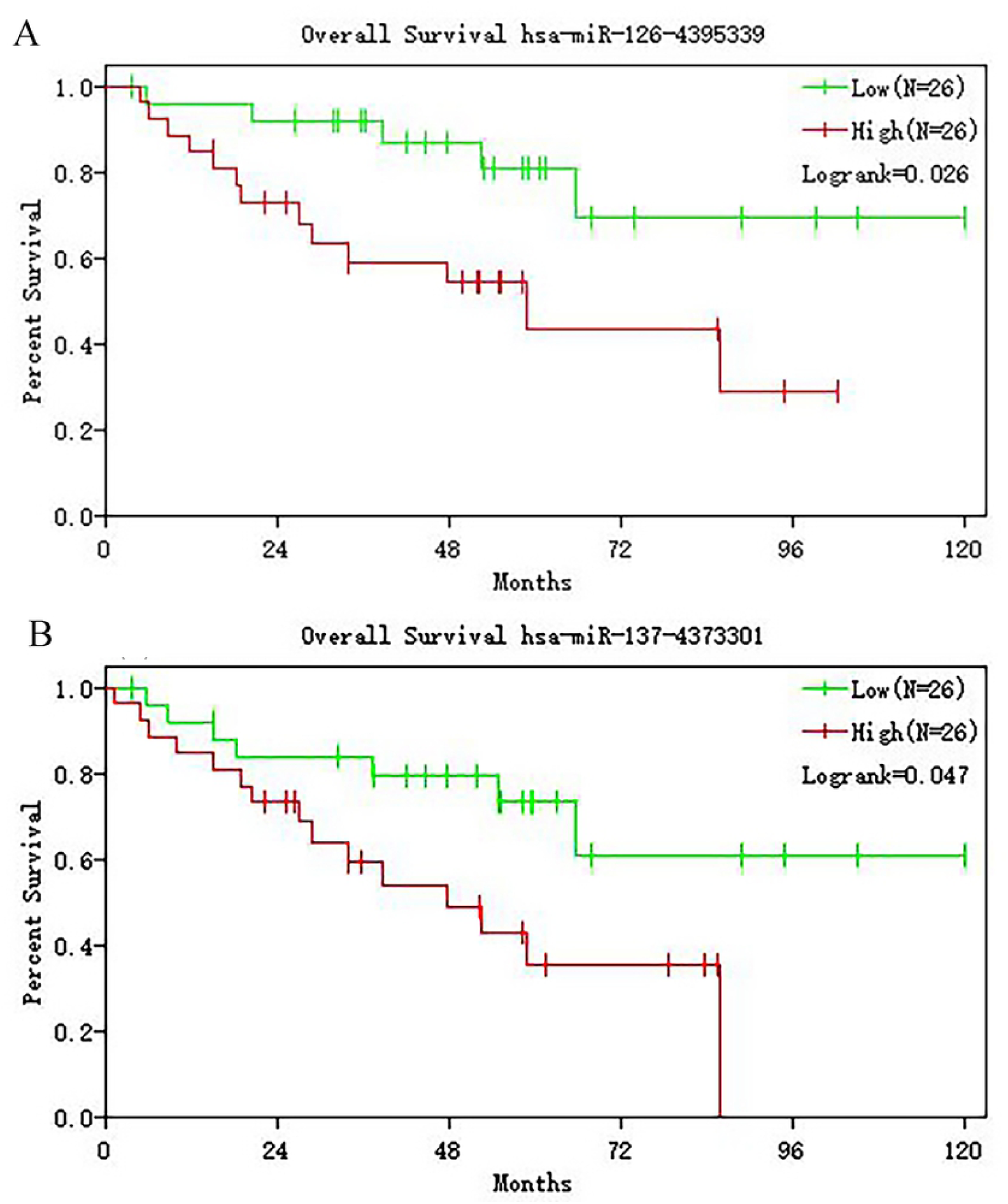
Network Nine. HUB-miRNAs in the center of the network were screened out.

**Figure 7.**
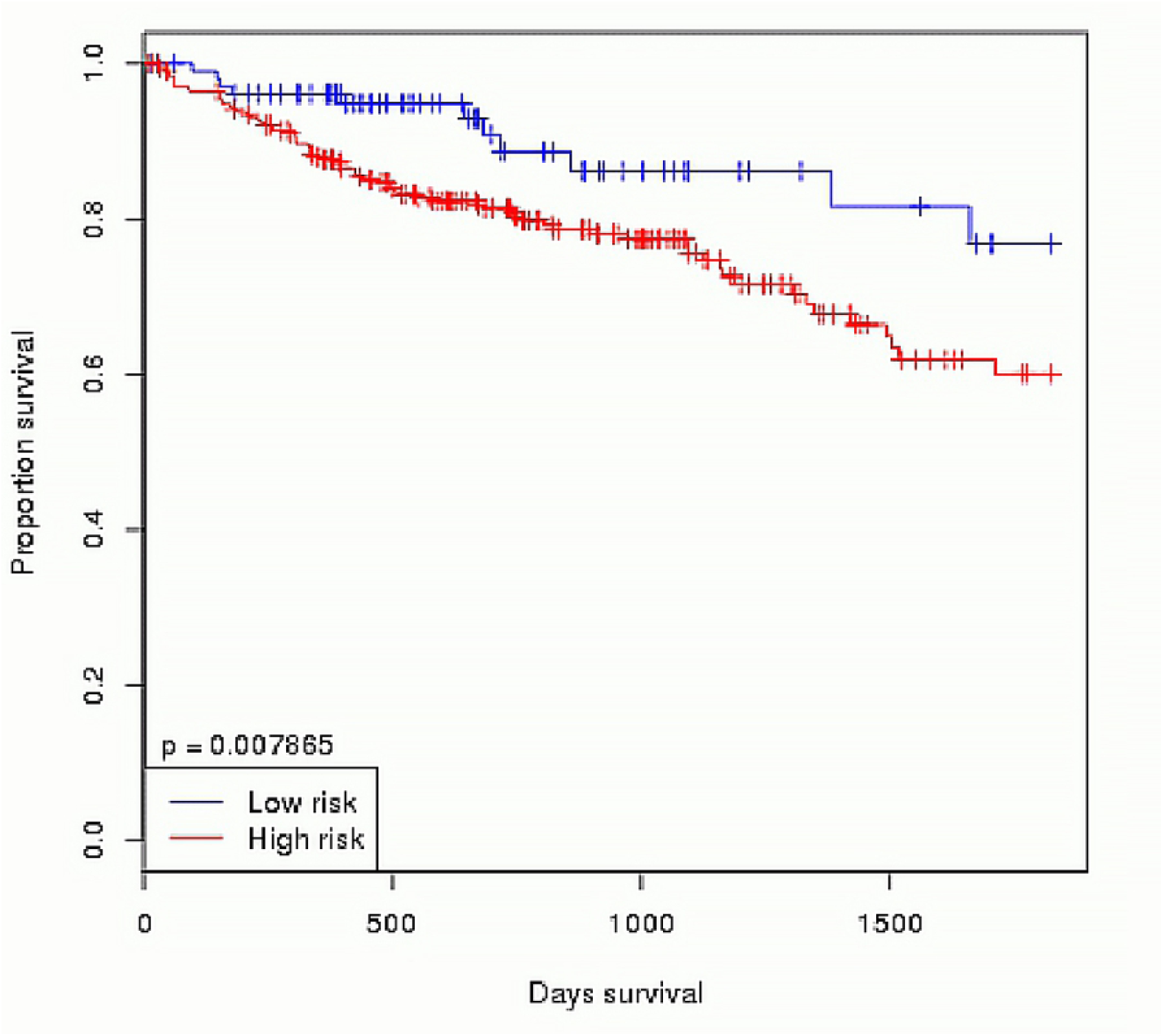
KEGG and reactome. (A) KEGG pathway analysis of miRNAs in turquoise module. (B) Reactome analysis of miRNAs in turquoise module.

**Figure 8.**
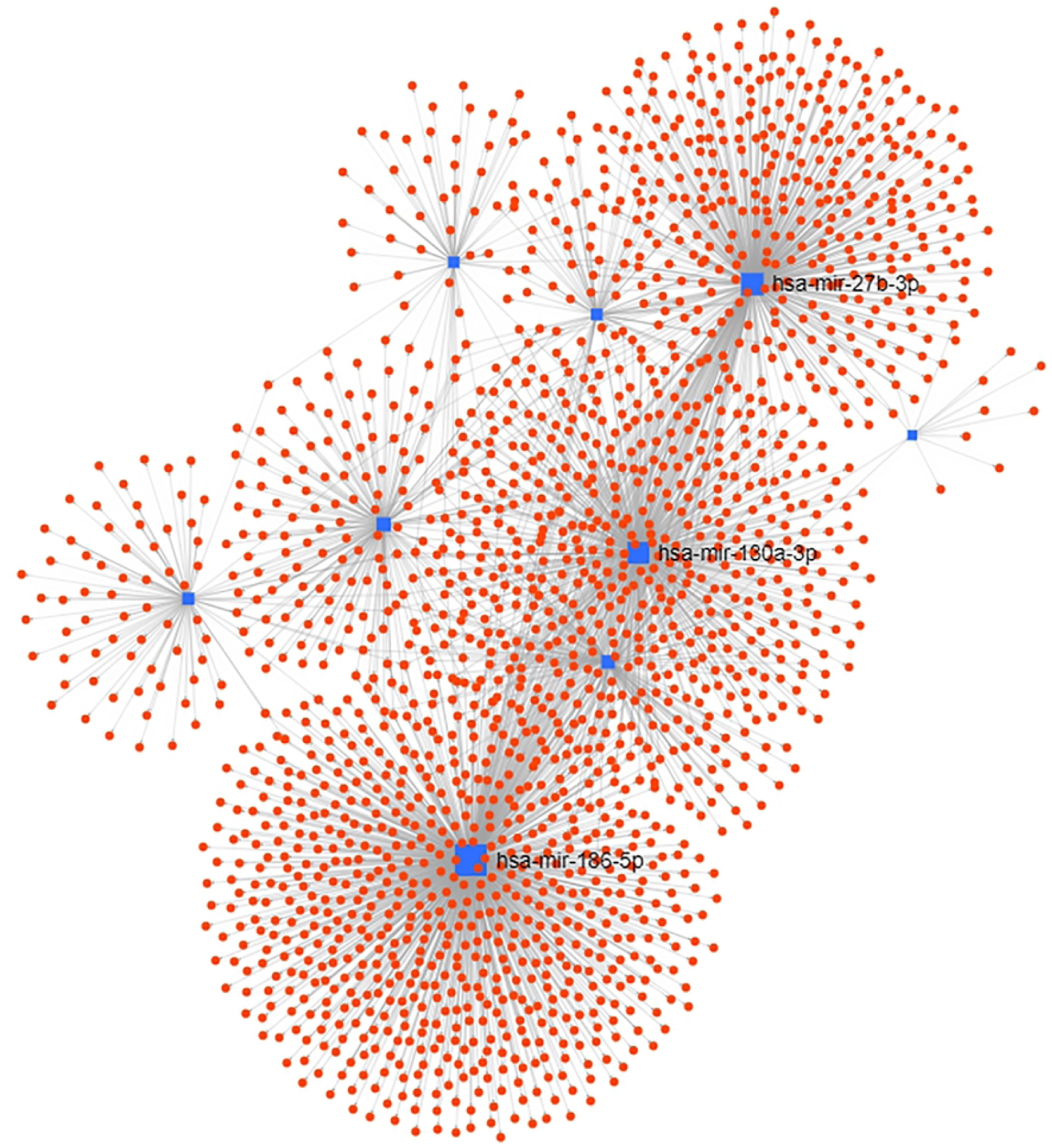
GO-MF. Molecular function analysis of miRNAs in turquoise module.

**Figure 9.**
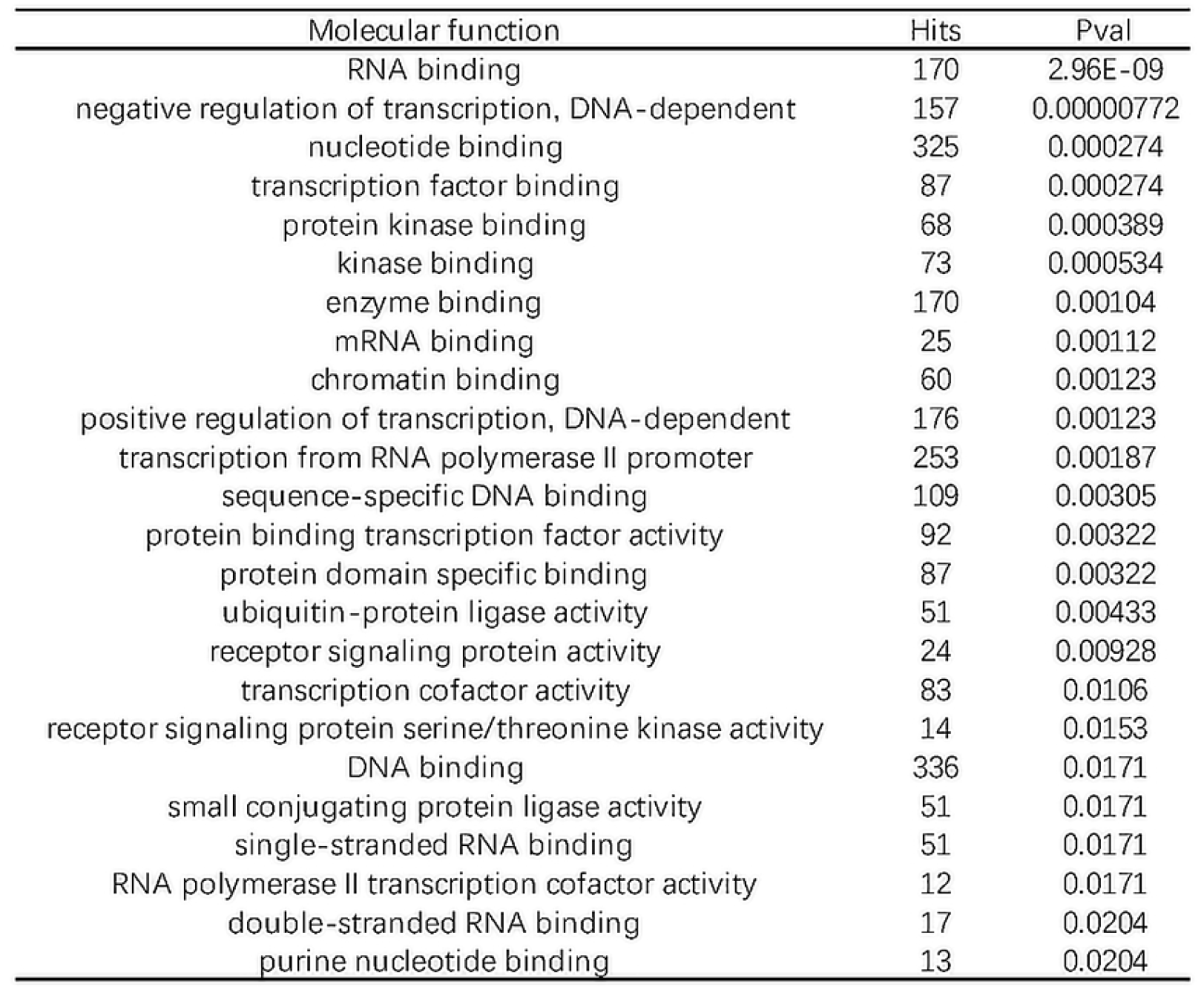
GO-CC. Cellular component analysis of miRNAs in turquoise module.

**Figure 10.**
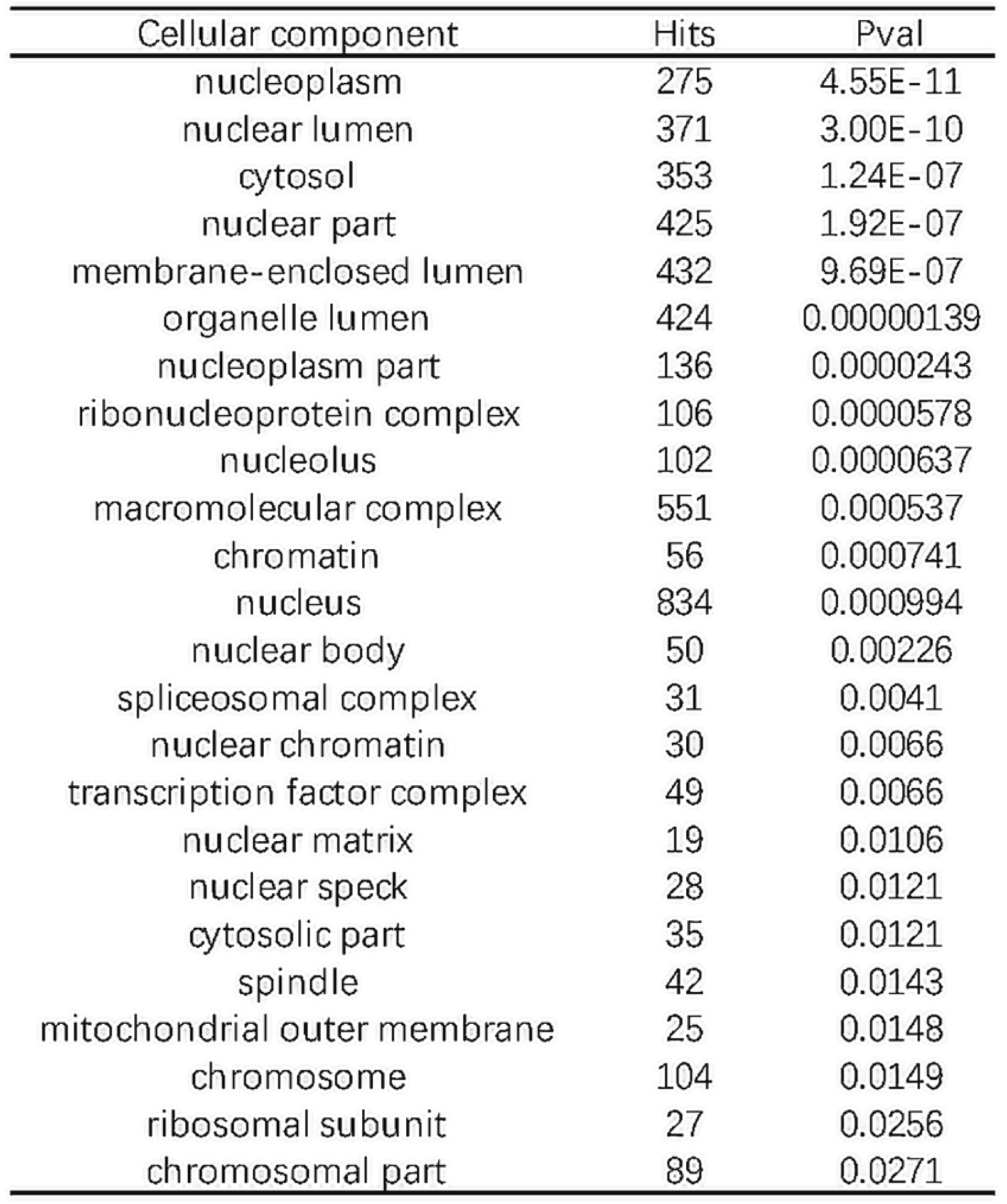
GO-BP. Biological process analysis of miRNAs in turquoise module.

### 3.3 Prognostic value verification

In order to validate the prognostic value of HUB-miRNAs found in this study as biomarkers, we evaluated whether HUB-miRNAs were associated with the overall survival rate of COAD patients. Cox regression and KM analysis were performed using GSE26923 (n=65) chips from GEO database. It was found that in the HUB-miRNA of turquoise hsa-mir-137 (P < 0.031636223) and hsa-mir-126-3p (P < 0.027019656) were significantly correlated with overall survival rate (Fig 11). Using ONCOMIR for survival analysis, it was found that has-miR-126-3p and has-miR-137 could divide patients with COAD into high-risk group and low-risk group S = 3.378*EmiR-137+1.473*EmiR-126-3p. P< 0.007865 (Fig 12).

**Figure 11.**
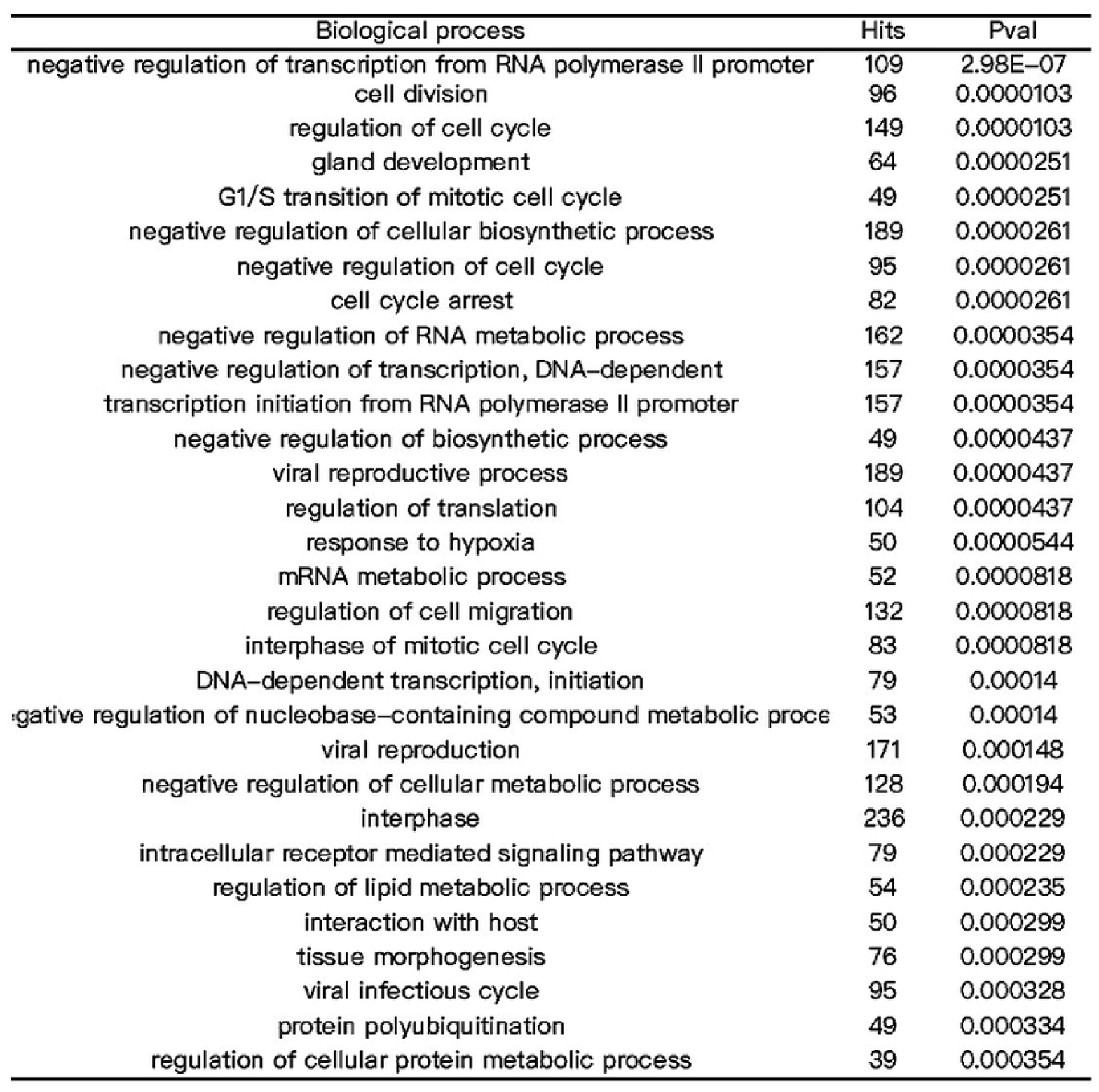
Overall survival rate of 2 HUB-miRNAs. (A) Hsa-mir-137 were significantly correlated with overall survival rate (P < 0.031636223). (B) Hsa-mir-126-3p were significantly correlated with overall survival rate (P < 0.027019656).

**Figure 12.**
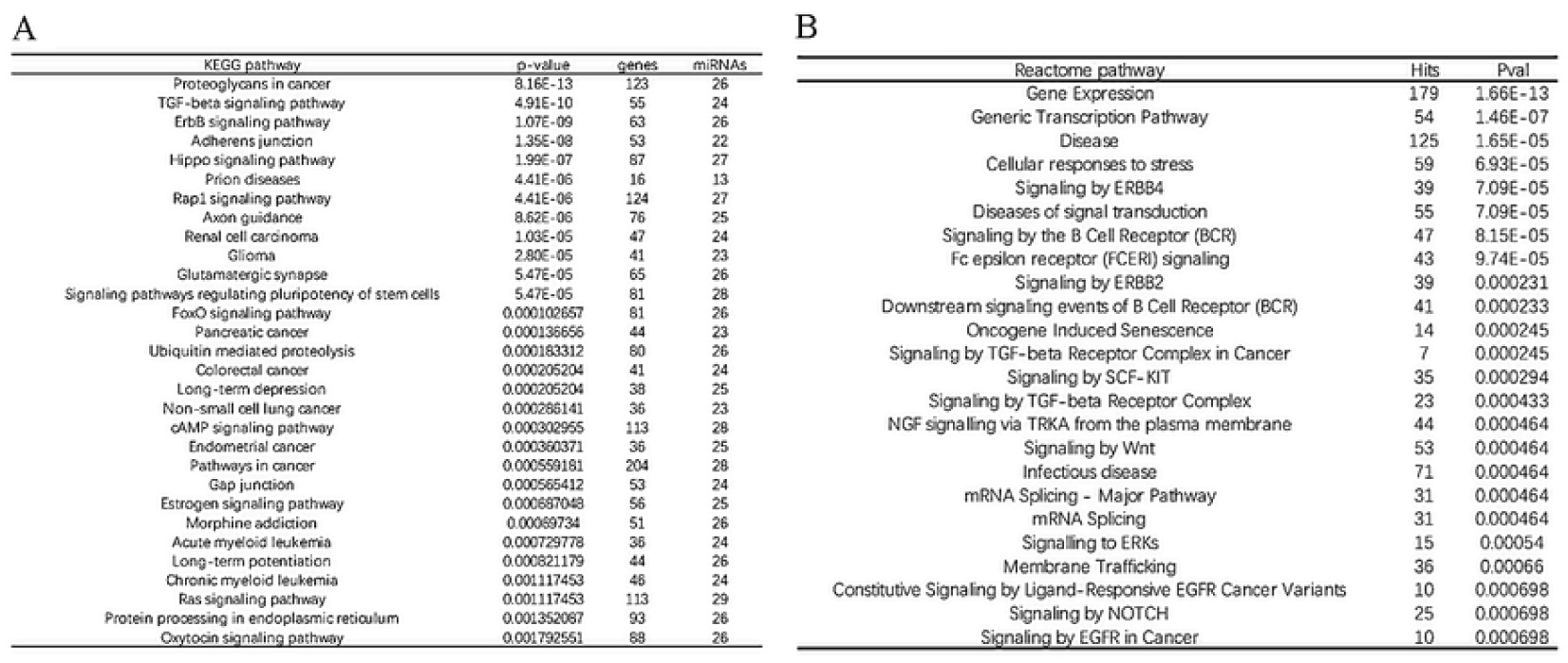
ONCOMIR validation. The combination of has-miR-126-3p and has-miR-137 could divide patients with COAD into high risk group and low risk group S = 3.378*EmiR-137+1.473*EmiR-126-3p. P< 0.007865

## 2 Discussion

Several cases of massive data mining miRNA with the WGCNA method have been reported. The characteristic of this method is that the correlation coefficient of the expression value is taken to the N power, so that their distribution is more in line with the scale-free network the true distribution in nature. It is used to describe the correlation pattern between genes with large amounts of data, full utilization of information makes it an ideal tool for predicting biomarkers(15, 20). It is common to select the few miRNAs that satisfy the condition of e | kME | > = threshold (0.8) for follow-up analysis, but it may lead to information loss. In the current study we select all the miRNAs in the module for network analysis, reduce the information loss, and focus on the center miRNAs which are closest to the relevant pathway. The molecular function and cell function were annotated by GO, we used another 65 samples of GSE26923 to verify the prognostic value of HUB-miRNAs.

In general, miRNA can play a negative role in the regulation of mRNA expression by binding to the complementary sequences in the target mRNA 3’-UTR, resulting in translation inhibition and / or target degradation(21). In recent years, increasing evidence has proved that miRNAs could also upregulate gene expression in specific cell types and conditions with distinct transcripts and proteins(22). It has found that the role of miRNA depends on MiRISC, and the dominant role of animal miRISC binding to target mRNAs are considered to be transcriptional instability, the increase of a certain miRNA may represent the instability of many cancer-related factors(23). It is an ideal and efficient therapeutic target, which is closely related to the occurrence and development of the tumor. KEGG analysis of the 9 candidate miRNAs in the turquoise module and their target genes showed that they were significantly associated with colon cancer (P = 4.09e-7), and with many pathways in cancer, including WNT and MAPK signals, TGF-beta and p53 VEGF and it is also closely related to cell cycle pathways. Our data show that these miRNAs are key regulators of these core carcinogenic functional pathways in COAD.

In order to confirm the prognostic value of HUB-miRNAs, we verified two miRNAs (hsa-mir-137, hsa-mir-126-3p) through the data from other sources, through which patients can be divided into the high-risk group and low-risk group. The HUB-miRNAs regulate 21 target genes that have been shown to be associated with colon cancer. It is reported hsa-mir-137 upregulated 6.6-fold in the lymph node positive colon cancer group compared with the lymph node negative colon cancer group(24). Its downstream target genes include CDK6, CDC42. CDK6 plays an important role in the progress of G 1 phase and G 1 / S transition in the cell cycle(25). It can regulate the activity of tumor suppressor protein Rb by phosphorylation and has been shown to act as an anti-tumor factor in a variety of cancers such as breast cancer / liver cancer / colon cancer(26). CDC42 plays an important role in regulating cell cycle(27), CDC42 expression was associated with a longer survival rate in breast cancer (P < 0.025)(28). Overexpression of hsa-mir-126-3p can promote cell proliferation and migration by activating PI3K/AKT signaling pathway(29). Hsa-mir-126-3p is reported as a novel regulator of PVC-mediated vessel stability during tumor angiogenesis(30). The expression of hsa-mir-126-3p is increased in patients with lung adenocarcinoma. As a biomarker for the diagnosis of early lung cancer, hsa-mir-126-3p is highly sensitive and specific(31). Its target genes include E2F1 VEGFA FOXO3. E2F1 gene polymorphism is considered to be associated with lung cancer head and neck cancer and other cancers. VEGFA gene polymorphism has been confirmed to be associated with metastatic colorectal cancer bladder cancer, renal cell carcinoma, prostate cancer and other cancers(32-35). FOXO 3a plays an inhibitory role in cancer and is closely related to malignant tumors such as breast cancer, colon cancer, prostate cancer, bladder cancer and nasopharyngeal carcinoma(36). We believe that these two kinds of miRNAs participate in the progress of COAD and increase the metastasis potential through their respective mechanisms. Their combination can be used as a biomarker to predict the behavior of COAD. It is a promising drug target for inhibiting tumor metastasis and increase survival rate.

There are also many imperfections in our research. The sample size of GSE chips for verification (n=65) is relatively smaller than WGCNA data (n=442). Fewer miRNAs were detected in GSE chips. There may be more HUB-miRNAs that have prognostic value. The direct experimental verification of the mechanism of HUB-miRNAs in the metastasis of colorectal adenocarcinoma needs to be done in the follow-up study.

This work was supported by the National Natural Science Foundation of China (http://www.nsfc.gov.cn/) under Grant No.81572082; Jilin Science and Technology Foundation (http://kjt.jl.gov.cn/) under Grant No.20150414015GH. The funders had no role in study design, data collection and analysis, decision to publish, or preparation of the manuscript.

## Disclosure of interest

The authors report no conflict of interest.

The datasets generated and analyzed in this paper are available in the TCGA (https://portal.gdc.cancer.gov/) and GEO (www.ncbi.nlm.nih.gov/geo/) databases.

## References

1. The L. Toward better control of colorectal cancer. The Lancet. 2014;383(9927). doi: 10.1016/s0140-6736(14)60699-1.

2. Kuipers EJ, Grady WM, Lieberman D, Seufferlein T, Sung JJ, Boelens PG, et al. Colorectal cancer. Nature reviews Disease primers. 2015;1:15065. Epub 2015/01/01. doi: 10.1038/nrdp.2015.65. PubMed PMID: 27189416; PubMed Central PMCID: PMCPMC4874655.

3. Bhat SK, East JE. Colorectal cancer: prevention and early diagnosis. Medicine. 2015;43(6):295–8. doi: 10.1016/j.mpmed.2015.03.009.

4. Favoriti P, Carbone G, Greco M, Pirozzi F, Pirozzi RE, Corcione F. Worldwide burden of colorectal cancer: a review. Updates in surgery. 2016;68(1):7–11. Epub 2016/04/14. doi: 10.1007/s13304-016-0359-y. PubMed PMID: 27067591.

5. Maida M, Macaluso FS, Ianiro G, Mangiola F, Sinagra E, Hold G, et al. Screening of colorectal cancer: present and future. Expert review of anticancer therapy. 2017;17(12):1131–46. Epub 2017/10/13. doi: 10.1080/14737140.2017.1392243. PubMed PMID: 29022408.

6. Moreno-Moya JM, Vilella F, Simon C. MicroRNA: key gene expression regulators. Fertility and sterility. 2014;101(6):1516–23. Epub 2013/12/10. doi: 10.1016/j.fertnstert.2013.10.042. PubMed PMID: 24314918.

7. Kontomanolis EN, Koukouli A, Liberis G, Stanulov H, Achouhan A, Pagkalos A. MiRNAs: regulators of human disease. European journal of gynaecological oncology. 2016;37(6):759–65. Epub 2016/01/01. PubMed PMID: 29943916.

8. Garofalo M, Croce CM. microRNAs: Master Regulators as Potential Therapeutics in Cancer. 2011;51(1):25–43. doi: 10.1146/annurev-pharmtox-010510-100517. PubMed PMID: 20809797.

9. Mishra S, Yadav T, Rani V. Exploring miRNA based approaches in cancer diagnostics and therapeutics. Critical reviews in oncology/hematology. 2016;98:12–23. Epub 2015/10/21. doi: 10.1016/j.critrevonc.2015.10.003. PubMed PMID: 26481951.

10. Thomas J, Ohtsuka M, Pichler M, Ling H. MicroRNAs: Clinical Relevance in Colorectal Cancer. International journal of molecular sciences. 2015;16(12):28063–76. Epub 2015/11/26. doi: 10.3390/ijms161226080. PubMed PMID: 26602923; PubMed Central PMCID: PMCPMC4691027.

11. Rupaimoole R, Slack FJ. MicroRNA therapeutics: towards a new era for the management of cancer and other diseases. Nature reviews Drug discovery. 2017;16(3):203–22. Epub 2017/02/18. doi: 10.1038/nrd.2016.246. PubMed PMID: 28209991.

12. Langfelder P, Horvath S. WGCNA: an R package for weighted correlation network analysis. BMC bioinformatics. 2008;9:559. Epub 2008/12/31. doi: 10.1186/1471-2105-9-559. PubMed PMID: 19114008; PubMed Central PMCID: PMCPMC2631488.

13. Zhang X, Feng H, Li Z, Li D, Liu S, Huang H, et al. Application of weighted gene co-expression network analysis to identify key modules and hub genes in oral squamous cell carcinoma tumorigenesis. Onco Targets Ther. 2018;11:6001–21. Epub 2018/10/03. doi: 10.2147/ott.S171791. PubMed PMID: 30275705; PubMed Central PMCID: PMCPMC6157991.

14. Su Q, Zhu EC, Qu YL, Wang DY, Qu WW, Zhang CG, et al. Serum level of co-expressed hub miRNAs as diagnostic and prognostic biomarkers for pancreatic ductal adenocarcinoma. Journal of Cancer. 2018;9(21):3991–9. Epub 2018/11/10. doi: 10.7150/jca.27697. PubMed PMID: 30410604; PubMed Central PMCID: PMCPMC6218787.

15. Yepes S, Lopez R, Andrade RE, Rodriguez-Urrego PA, Lopez-Kleine L, Torres MM. Co-expressed miRNAs in gastric adenocarcinoma. Genomics. 2016;108(2):93–101. Epub 2016/07/17. doi: 10.1016/j.ygeno.2016.07.002. PubMed PMID: 27422560.

16. Zhang B, Horvath S. A general framework for weighted gene co-expression network analysis. Statistical applications in genetics and molecular biology. 2005;4:Article17. Epub 2006/05/02. doi: 10.2202/1544-6115.1128. PubMed PMID: 16646834.

17. Fan Y, Habib M, Xia J. Xeno-miRNet: a comprehensive database and analytics platform to explore xeno-miRNAs and their potential targets. PeerJ. 2018;6:e5650. Epub 2018/10/04. doi: 10.7717/peerj.5650. PubMed PMID: 30280028; PubMed Central PMCID: PMCPMC6166626.

18. Vlachos IS, Zagganas K, Paraskevopoulou MD, Georgakilas G, Karagkouni D, Vergoulis T, et al. DIANA-miRPath v3.0: deciphering microRNA function with experimental support. Nucleic acids research. 2015;43(W1):W460–6. Epub 2015/05/16. doi: 10.1093/nar/gkv403. PubMed PMID: 25977294; PubMed Central PMCID: PMCPMC4489228.

19. Fan Y, Xia J. miRNet—Functional Analysis and Visual Exploration of miRNA–Target Interactions in a Network Context. In: von Stechow L, Santos Delgado A, editors. Computational Cell Biology: Methods and Protocols. New York, NY: Springer New York; 2018. p. 215–33.

20. Giulietti M, Occhipinti G, Principato G, Piva F. Identification of candidate miRNA biomarkers for pancreatic ductal adenocarcinoma by weighted gene co-expression network analysis. Cellular oncology (Dordrecht). 2017;40(2):181–92. Epub 2017/02/17. doi: 10.1007/s13402-017-0315-y. PubMed PMID: 28205147.

21. Towler BP, Jones CI, Newbury SF. Mechanisms of regulation of mature miRNAs. Biochemical Society transactions. 2015;43(6):1208–14. Epub 2015/11/29. doi: 10.1042/bst20150157. PubMed PMID: 26614662.

22. Valinezhad Orang A, Safaralizadeh R, Kazemzadeh-Bavili M. Mechanisms of miRNA-Mediated Gene Regulation from Common Downregulation to mRNA-Specific Upregulation. International journal of genomics. 2014;2014:970607. Epub 2014/09/03. doi: 10.1155/2014/970607. PubMed PMID: 25180174; PubMed Central PMCID: PMCPMC4142390.

23. Steinkraus BR, Toegel M, Fulga TA. Tiny giants of gene regulation: experimental strategies for microRNA functional studies. Wiley interdisciplinary reviews Developmental biology. 2016;5(3):311–62. Epub 2016/03/08. doi: 10.1002/wdev.223. PubMed PMID: 26950183; PubMed Central PMCID: PMCPMC4949569.

24. Huang ZM, Yang J, Shen XY, Zhang XY, Meng FS, Xu JT, et al. MicroRNA expression profile in non-cancerous colonic tissue associated with lymph node metastasis of colon cancer. Journal of digestive diseases. 2009;10(3):188–94. Epub 2009/08/08. doi: 10.1111/j.1751-2980.2009.00384.x. PubMed PMID: 19659786.

25. Enders GH. Mammalian interphase cdks: dispensable master regulators of the cell cycle. Genes & cancer. 2012;3(11-12):614–8. Epub 2013/05/02. doi: 10.1177/1947601913479799. PubMed PMID: 23634250; PubMed Central PMCID: PMCPMC3636753.

26. Bevara GB, Naveen Kumar AD, Koteshwaramma KL, Badana A, Kumari S, Malla RR. C-glycosyl flavone from Urginea indica inhibits proliferation & angiogenesis & induces apoptosis via cyclin-dependent kinase 6 in human breast, hepatic & colon cancer cell lines. The Indian journal of medical research. 2018;147(2):158–68. Epub 2018/05/29. doi: 10.4103/ijmr.IJMR_51_16. PubMed PMID: 29806604; PubMed Central PMCID: PMCPMC5991124.

27. Olson MF, Ashworth A, Hall A. An essential role for Rho, Rac, and Cdc42 GTPases in cell cycle progression through G1. Science (New York, NY). 1995;269(5228):1270–2. Epub 1995/09/01. PubMed PMID: 7652575.

28. Chrysanthou E, Gorringe KL, Joseph C, Craze M, Nolan CC, Diez-Rodriguez M, et al. Phenotypic characterisation of breast cancer: the role of CDC42. Breast cancer research and treatment. 2017;164(2):317–25. Epub 2017/04/30. doi: 10.1007/s10549-017-4267-8. PubMed PMID: 28451966; PubMed Central PMCID: PMCPMC5487723.

29. Chang L, Liang J, Xia X, Chen X. miRNA-126 enhances viability, colony formation, and migration of keratinocytes HaCaT cells by regulating PI3 K/AKT signaling pathway. Cell biology international. 2019;43(2):182–91. Epub 2018/12/21. doi: 10.1002/cbin.11088. PubMed PMID: 30571843.

30. Pitzler L, Auler M, Probst K, Frie C, Bergmeier V, Holzer T, et al. miR-126-3p Promotes Matrix-Dependent Perivascular Cell Attachment, Migration and Intercellular Interaction. Stem cells (Dayton, Ohio). 2016;34(5):1297–309. Epub 2016/03/05. doi: 10.1002/stem.2308. PubMed PMID: 26934179.

31. Kim JE, Eom JS, Kim WY, Jo EJ, Mok J, Lee K, et al. Diagnostic value of microRNAs derived from exosomes in bronchoalveolar lavage fluid of early-stage lung adenocarcinoma: A pilot study. Thoracic cancer. 2018;9(8):911–5. Epub 2018/05/29. doi: 10.1111/1759-7714.12756. PubMed PMID: 29806739; PubMed Central PMCID: PMCPMC6068458.

32. Bendardaf R, Sharif-Askari FS, Sharif-Askari NS, Syrjanen K, Pyrhonen S. Cytoplasmic E-Cadherin Expression Is Associated With Higher Tumour Level of VEGFA, Lower Response Rate to Irinotecan-based Treatment and Poorer Prognosis in Patients With Metastatic Colorectal Cancer. Anticancer research. 2019;39(4):1953–7. Epub 2019/04/07. doi: 10.21873/anticanres.13305. PubMed PMID: 30952738.

33. Song Y, Yang Y, Liu L, Liu X. Association between five polymorphisms in vascular endothelial growth factor gene and urinary bladder cancer risk: A systematic review and meta-analysis involving 6671 subjects. Gene. 2019;698:186–97. Epub 2019/03/09. doi: 10.1016/j.gene.2019.02.070. PubMed PMID: 30849545.

34. Wang J, Shen C, Fu Y, Yu T, Song J. The associations between five polymorphisms of vascular endothelial growth factor and renal cell carcinoma risk: an updated meta-analysis. Onco Targets Ther. 2017;10:1725–34. Epub 2017/03/31. doi: 10.2147/ott.S125965. PubMed PMID: 28356760; PubMed Central PMCID: PMCPMC5367456.

35. Song Y, Hu J, Chen Q, Guo J, Zou Y, Zhang W, et al. Association between vascular endothelial growth factor rs699947 polymorphism and the risk of three major urologic neoplasms (bladder cancer, prostate cancer, and renal cell carcinoma): A meta-analysis involving 11,204 subjects. Gene. 2018;679:241–52. Epub 2018/09/10. doi: 10.1016/j.gene.2018.09.005. PubMed PMID: 30195633.

36. Liu Y, Ao X, Ding W, Ponnusamy M, Wu W, Hao X, et al. Critical role of FOXO3a in carcinogenesis. Molecular cancer. 2018;17(1):104. Epub 2018/07/27. doi: 10.1186/s12943-018-0856-3. PubMed PMID: 30045773; PubMed Central PMCID: PMCPMC6060507.

